## Supplementary Information for "A Versatile Soluble Siglec Scaffold for Sensitive and Quantitative Detection of Glycan Ligands"

**Supplementary Table 1:** Siglec-Fc expression levels CHO WT and CHO Lec1 cells as determined by ELISA. n.d. = not determined.

| Siglec | CHO WT (mg/L) | CHO Lec1 (mg/L) |
|---|---|---|
| 1 | 8.8 | 5.4 |
| 2 (V1) | 10 | N/A |
| 2 (V2) | 19 | 15 |
| 3 | 5.4 | 12 |
| 4 | 6.5 | 14 |
| 5 | 36 | 5.4 |
| 6 | 39 | 43 |
| 7 | 20 | 12 |
| 8 | 6.7 | 10 |
| 9 | 15 | 17 |
| 10 | 8.4 | 6.7 |
| 11 | 7.1 | 8.1 |
| 12 | 25 | n.d. |
| 14 | 42 | n.d. |
| 15 | 11 | n.d. |
| 16 | 8.4 | n.d. |

**Supplementary Table 2: Primers for developing the Siglec-Fcs in the 2-domain and 3-domain variants.** GCTAGC is the restriction sequence for NheI used in the forward primer to generate the 5’ restriction site. ACCGGT is the restriction sequence for AgeI used in the reverse primers to generate the 3’ restriction site. The AGCAGC leader sequence is used to stabilize the primers and assist the restriction enzyme during digestion.

| Human Siglec | Forward primer | Reverse primer |
|---|---|---|
| Siglec-1 | agcagcgctagcatgggcttcttgcccaagcttc | agcagcaccggtctggacctcagccatgaagatg |
| CD22 | agcagcgctagcatgcatctcctcggcccctgg | agcagcaccggtcgtggaaggttccggggcatac |
| CD33 | agcagcgctagcatgccgctgctgctactgctg | agcagcaccggtatgaaccactcctgctctgg |
| Siglec-4 | agcagcgctagcatgatattcctcacggcactg | agcagcaccggttgtcccgttcactgttggcttc |
| Siglec-5 | agcagcgctagcatgctgcccctgctgctgctg | agcagcaccggtggggcccagcaactgtgggag |
| Siglec-6 | agcagcgctagcatgcagggagcccaggaagcc | agcagcaccggtcctgccttctggtttccaatg |
| Siglec-7 | agcagcgctagcatgctgctgctgctgctgctg | agcagcaccggtcctcattttgcctgtgtactc |
| Siglec-8 | agcagcgctagcatgctgctgctgctgctgctg | agcagcaccggttcttgaggtgcctgtgccctc |
| Siglec-9 | agcagcgctagcatgctgctgctgctgctgcccc | agcagcaccggtagtcactcctgatgtggctttg |
| Siglec-10 | agcagcgctagcatgctactgccactgctgctg | agcagcaccggttctcaggttctctggaggatac |
| Siglec-11 | agcagcgctagcatgctgctgctgcccctgctg | agcagcaccggtgttttccaggactgtcctgtttg |
| Siglec-12 | agcagcgctagcatgctactgctgctgctactg | agcagcaccggtgatggagaaggtgggcatgtg |
| Siglec-14 | agcagcgctagcatgctgcccctgctgctgctg | agcagcaccggtagaggagcttctctgcacag |
| Siglec-15 | agcagcgctagcatggaaaagtccatctggctg | agcagcaccggtggccccgctggcgccatggaag |
| Siglec-16 | agcagcgctagc atgctgctgctgcccctgctg | agcagcaccggtcactctcaggttctctggagg |

**Supplementary Table 3: Mutagenesis primers for developing arginine mutant human Siglecs.** The bolded bases represent the mutation codon to an alanine from the original arginine. The specific arginine residue mutated to an alanine for each Siglec is outlined in the second column.

| Human Siglec | Forward primer | Reverse primer |
|---|---|---|
| Siglec-1 | ctctggttcctacaacttc**gcc**ttcgaaatcagtgaggtc | gacctcactgatttcgaa**ggc**gaagttgtaggaaccagag |
| CD22 | gtggtcagctggggctg**gcg**atggagtccaagactgag | ctcagtcttggactccat**cgc**cagccccagctgaccac |
| CD33 | gataatggttcatacttcttt**gcg**atggagagaggaagtacc | ggtacttcctctctccat**cgc**aaagaagtatgaaccattatc |
| Siglec-4 | ggcgggaagtactacttc**gct**ggggacctgggcggctac | gtagccgcccaggtcccc**agc**gaagtagtacttcccgcc |
| Siglec-5 | cttccgcgtggagagagga**gcg**gatgtaaaatatagctac | gtagctatattttacatc**cgc**tcctctctccacgcggaag |
| Siglec-6 | caatgctgcatacttcttt**gcg**ttgaagtccaaatggatg | catccatttggacttcaa**cgc**aaagaagtatgcagcattg |
| Siglec-7 | caaaaattgcaccctgagcatc**gca**gatgccagaatgagtgatg | catcactcattctggcatc**tgc**gatgctcagggtgcaatttttg |
| Siglec-8 | catatttctttcggctagag**gca**ggaagcatgaaatggagttac | gtaactccatttcatgcttcc**tgc**ctctagccgaaagaaatatg |
| Siglec-9 | gcggggagatacttctttgctatggagaaaggaagtataaaatg | catttatacttcctttctccatagcaaagaagtatctccccgc |
| Siglec-10 | gagtcacagtacttcttt**gcg**gtggagagaggaagctatg | catagcttcctctctccac**cgc**aaagaagtactgtgactc |
| Siglec-11 | gaggcatggtacttcttt**gcg**gtggagagaggaagccgtg | cacggcttcctctctccaccgcaaagaagtaccatgcctc |
| Siglec-12 | N/A | N/A |
| Siglec-14 | cacgggaagctatttcttc**gcc**gtggagagaggaagggatg | catcccttcctctctccac**ggc**gaagaaatagcttcccgtg |
| Siglec-15 | gaccgccgctacttctgcgccgtcgagttcgccggcgac | gtcgccggcgaactcgacggcgcagaagtagcggcggtc |
| Siglec-16 | gaggcatggtacttcttt**gcg**gtggagagaggaagccgtg | cacggcttcctctctccac**cgc**aaagaagtaccatgcctc |

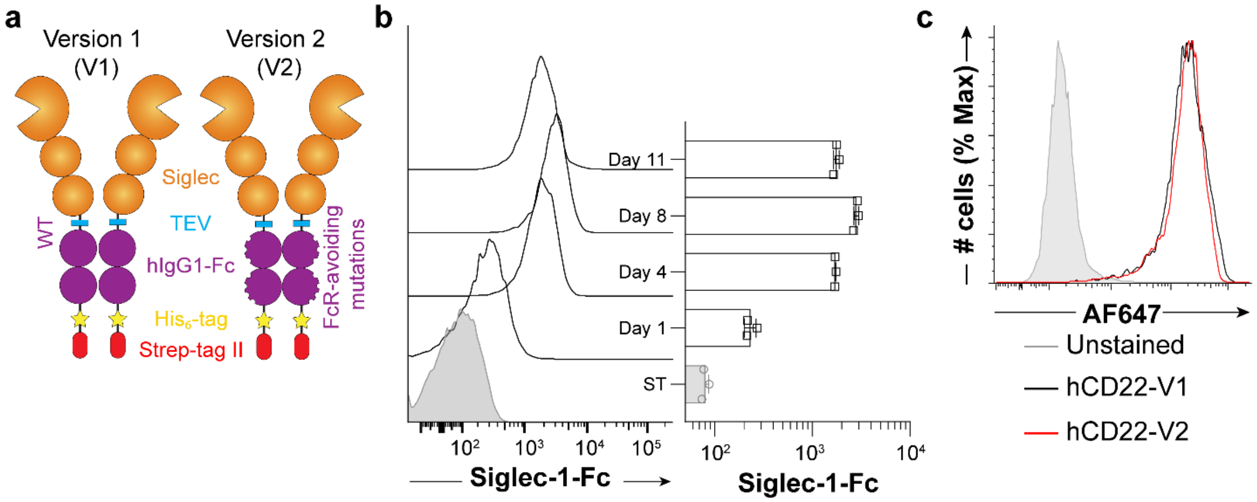


**Supplementary Figure 1: Validation of new constructs binding to cells. a**, Depiction of features built into the new Siglec-Fc constructs. The key difference between Version 1 (V1) and Version 2 (V2) are point mutants in the Fc that prevent binding to FcγRs. **b**, Expression levels of CD22-Fc as days post confluency as determined by ELISA and flow cytometry analysis of binding to U937 cells. **c**, Human CD22-Fc in either V1 (Black) or V2 (Red) constructs binding to ST6Gal1-overexpressing CHO cells, detected by anti-human IgG1-AF647 secondary antibody. Error bars represent +/- standard deviation of three replicates.

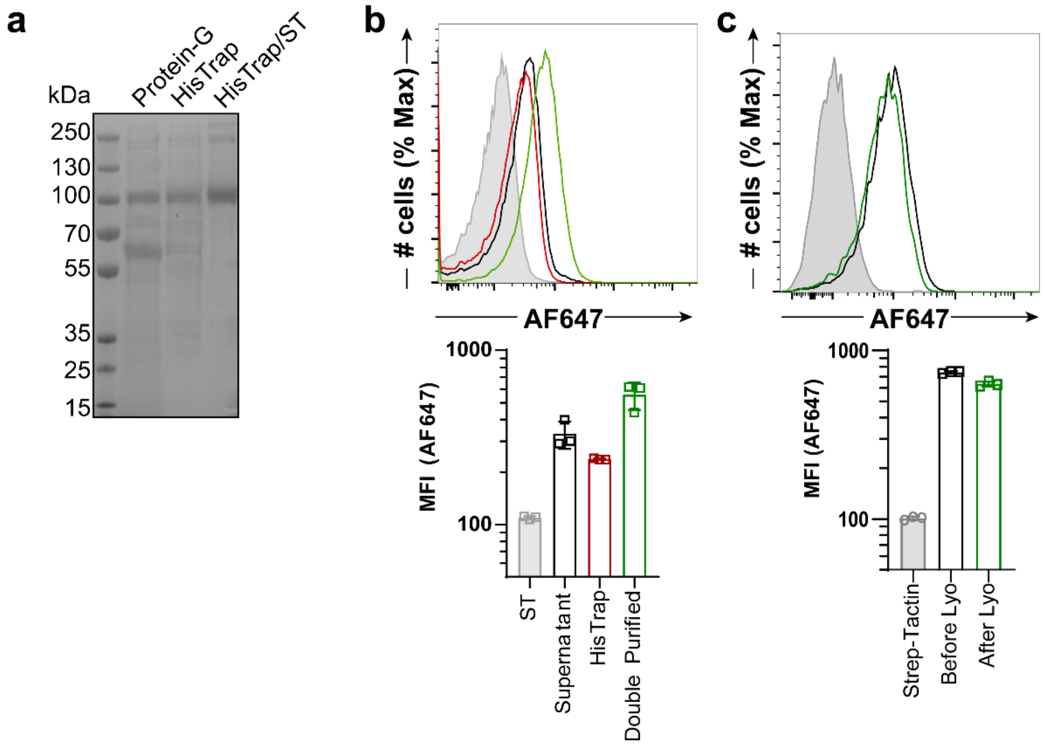


**Supplementary Figure 2: Testing binding after purification and lyophilizaiton. a**, SDS-PAGE gel of Sig-6-Fc purified using three purification strategies (ST; Strep-Tactin). **b**, Binding of Siglec-6-Fc to HEK293T cells either from supernatant directly (Black), purified through HisTrap only (Red), or double purified through HisTrap and Strep-Tactin column (Green). **c**, Binding of Siglec-6-Fc to HEK293T cells before or after lyophilization of aliquots. Error bars represent +/- standard deviation of three replicates.

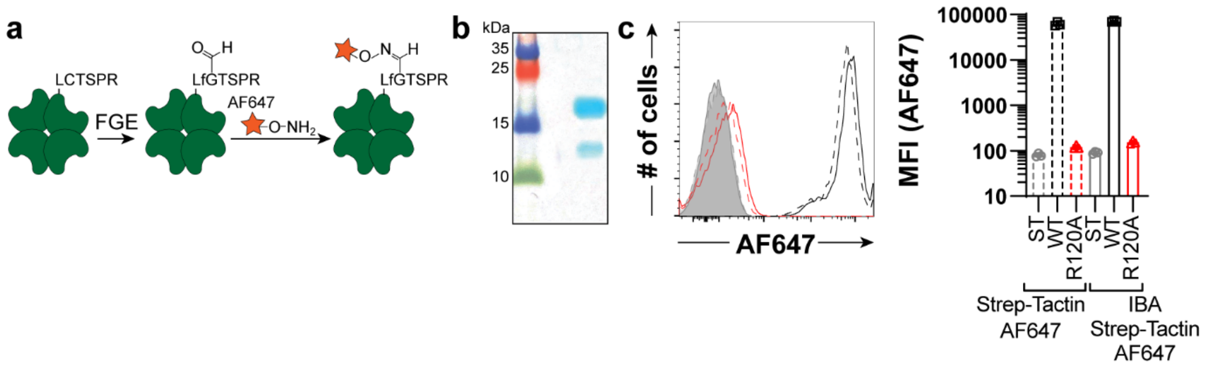


**Supplementary Figure 3: Creation and validation of Strep-Tactin-AF647. a**, Scheme of labelling Strep-Tactin with AF647 through the FGE consensus site. **b**, SDS-PAGE of labelled Strep-Tactin-AF647. **c**, Flow cytometry data of Siglec-1-Fc WT (Black) or Siglec-1-Fc R120A (Red) pre-complexed with Strep-Tactin-AF647 (IBA) (Solid lines) or our own Strep-Tactin-AF647 made (Dashed lines). Error bars represent +/- standard deviation of three replicates.

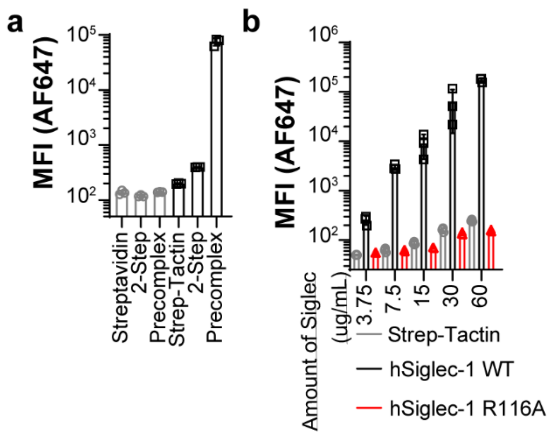


**Supplementary Figure 4: Increase in sensitivity of Strep-Tactin-AF647 over Streptavidin. a**, Binding of Siglec-1-Fc to U937 cells by pre-complexing with either Streptavidin (Grey bars) or Strep-Tactin-AF647 (Black bars). **b**, Titrating the amount of Siglec-1-Fc pre-complexed together and then incubated with U937 cells. Error bars represent +/- standard deviation of three replicates.

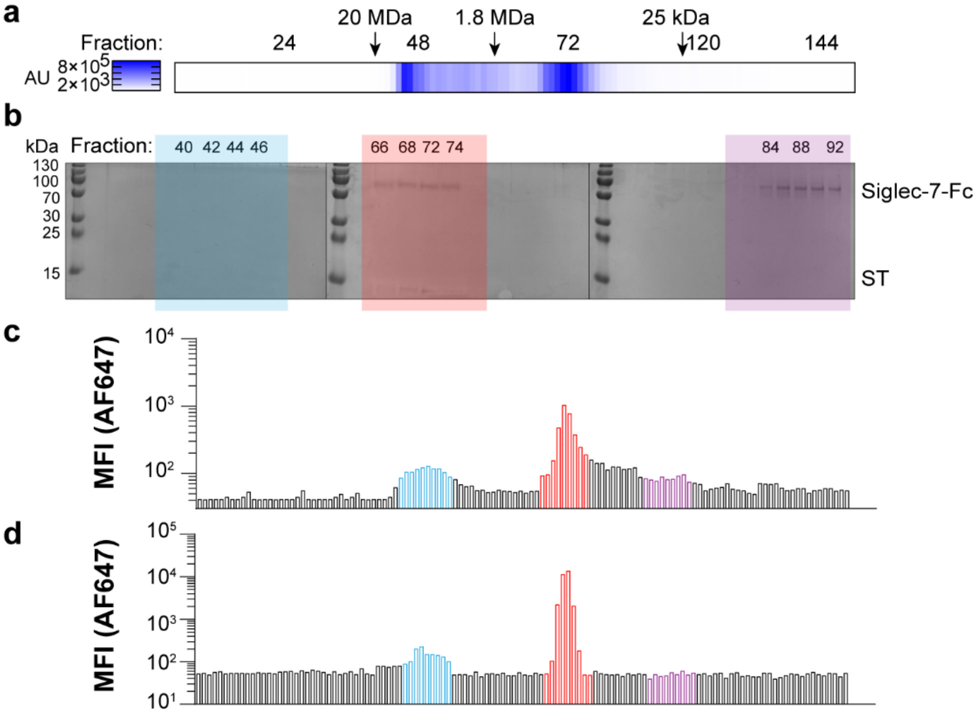


**Supplementary Figure 5: Validation of size of Siglec/Strep-Tactin-AF647 complex**. **a**, Siglec-7-Fc pre-complexed with Strep-Tactin-AF647 was passed over a gel filtration column and AF647 signal in the fraction. AU = arbitrary unity, which is the fluorescence of AF647. **b**, SDS-PAGE of the fractions (**c**), and binding of fractions to K562 cells by flow cytometry was carried out. **d,** The Siglec-7-Fc-Strep-Tactin-AF647 complex was lyophilized, resuspend, and tested for binding to K562 cells before and after lyophilization.

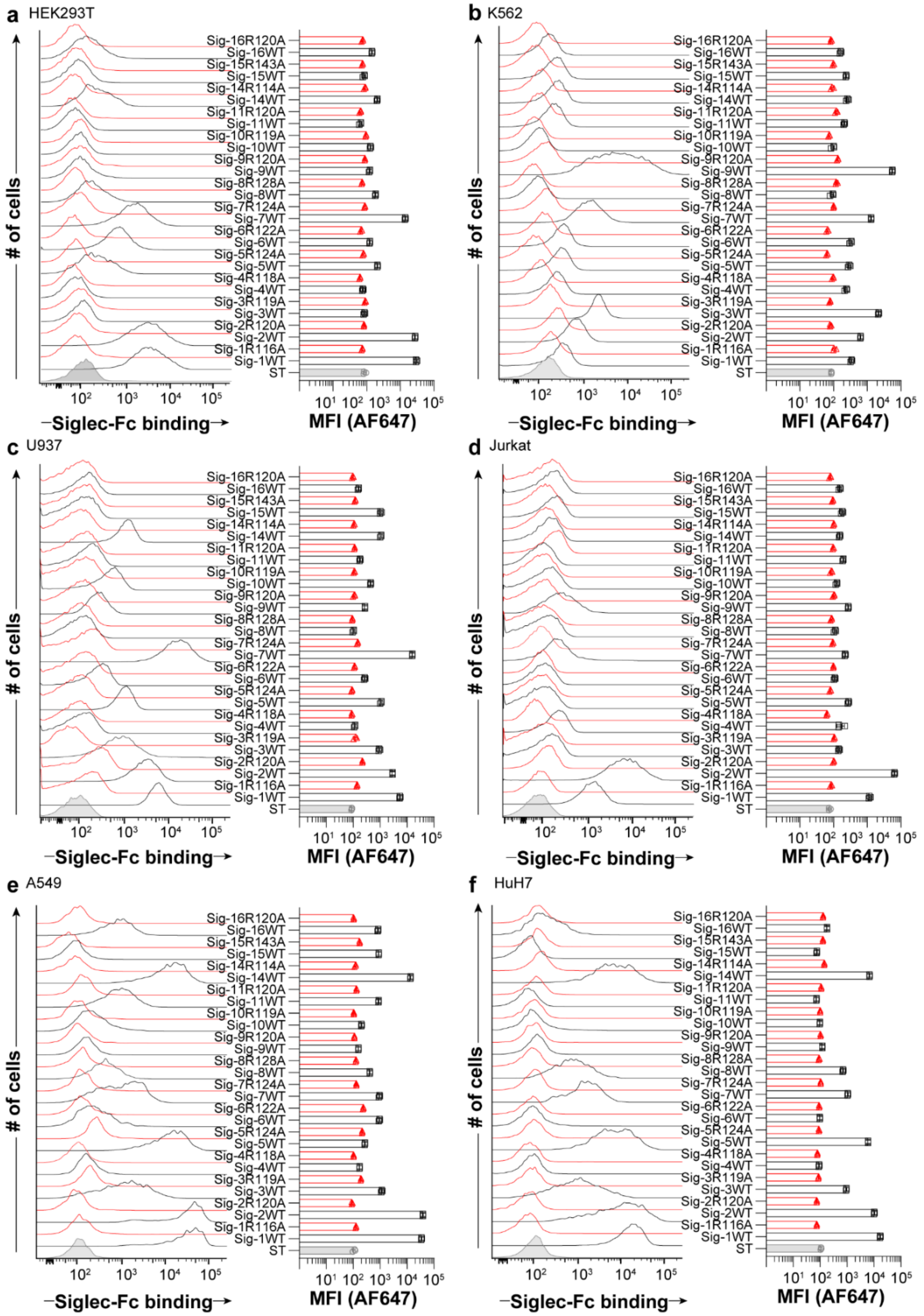


**Supplementary Figure 6: Probing Siglec ligands on cell lines.** Flow cytometry data of Siglec-Fc constructs pre-complexed with Strep-Tactin-AF647 binding to: **a**, HEK293T; **b**, K562; **c** U937; **d**, Jurkat; **e**, A549, and **f**, HuH7 cells. Error bars represent +/- standard deviation of three replicates.

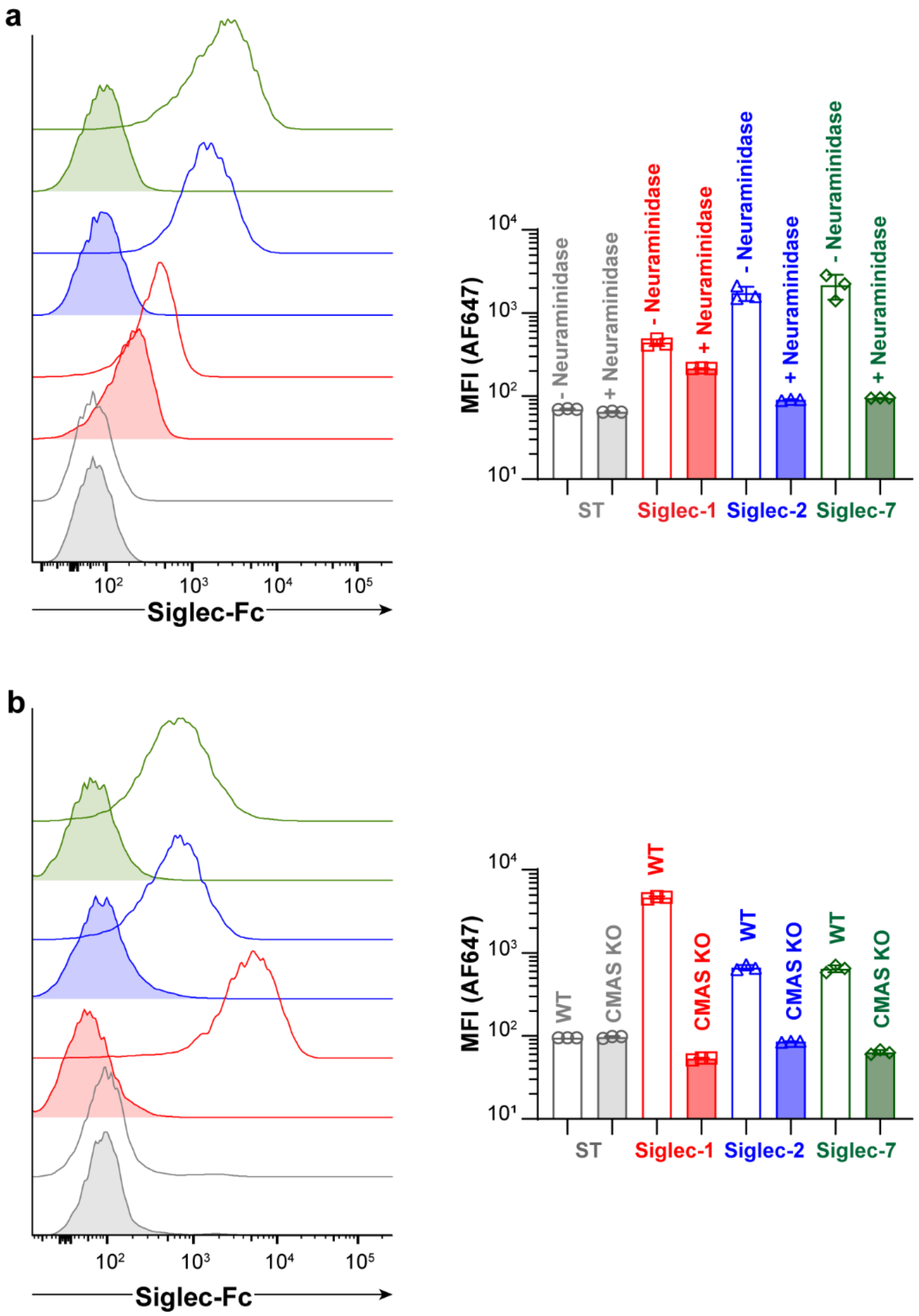


**Supplementary Figure 7: Validation of sialic acid dependent binding using the Siglec-Fc proteins. a**, Flow cytometry plots of Siglec-1, -2, and -7 WT pre-complexed with Strep-Tactin-AF647 and then incubated with K562 cells with or with pretreatment with Neu-A. **b**, Flow cytometry plots of Siglec-1, -2, and -7 (WT and corresponding essential arginine mutant) pre-complexed with Strep-Tactin-AF647 and then incubated with HEK293 WT or HEK293 CMAS KO cells. Error bars represent +/- standard deviation of three replicates.

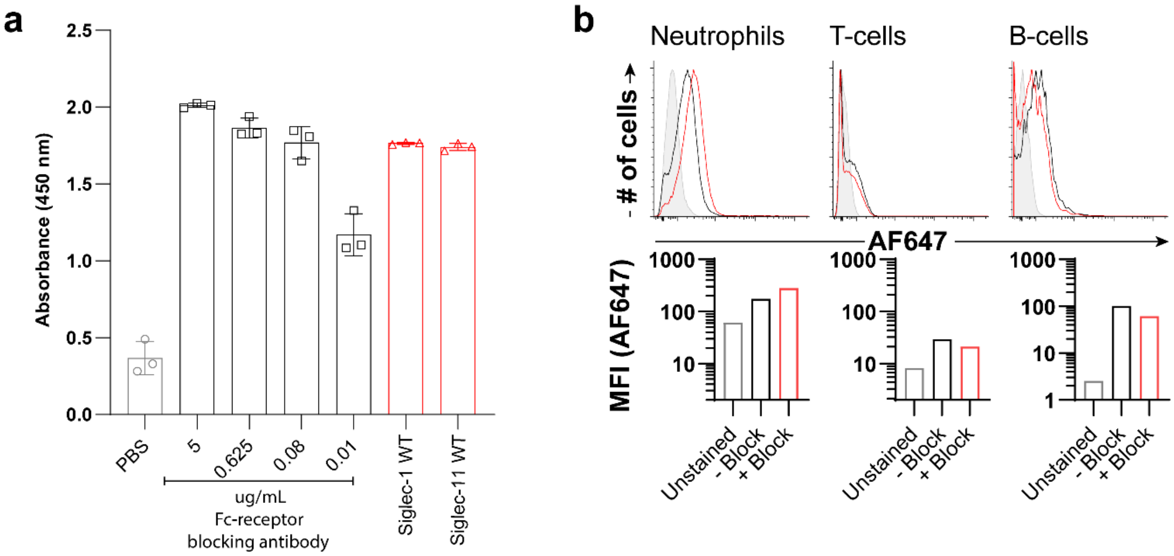


**Supplementary Figure 8: Binding of αhIgG1 to Fc-receptor blocking antibodies. a,** Direct ELISA of varying concentrations of Fc-receptor blocking antibody detected by commercial αhIgG1 secondary antibody. Positive control is Siglec-1 WT and Siglec-11 WT both plated at 9.6 µg/mL concentration. **b**, Flow cytometry data of αhIgG1 binding to Neutrophils, T-cells, and B-cells from human peripheral blood either pretreated with Fc receptor blocking antibodies (red) or without (black). Error bars represent +/- standard deviation of three replicates.

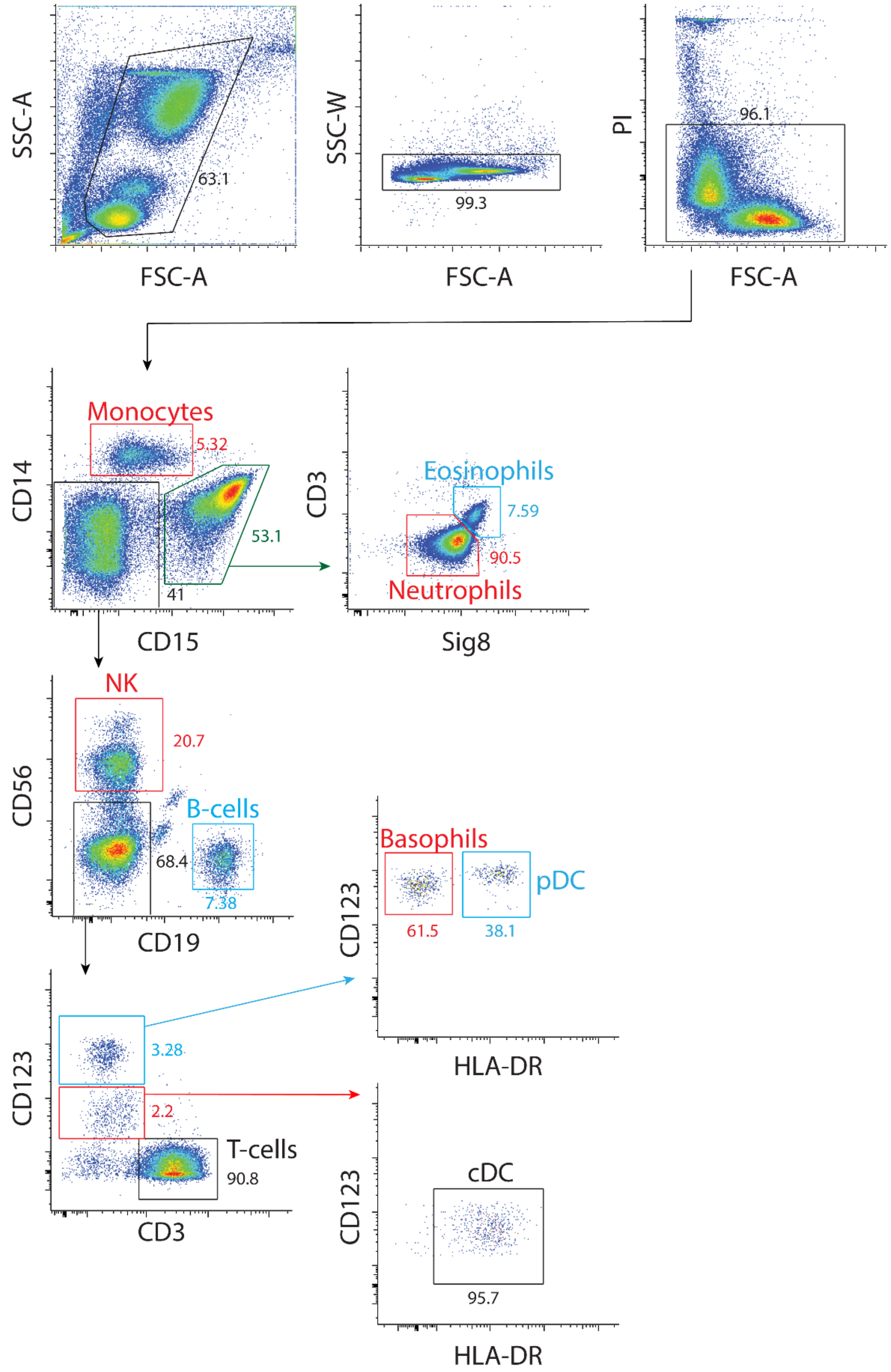


**Supplementary Figure 9: Representative gating strategy for analysis of human splenocytes and human peripheral blood samples.**

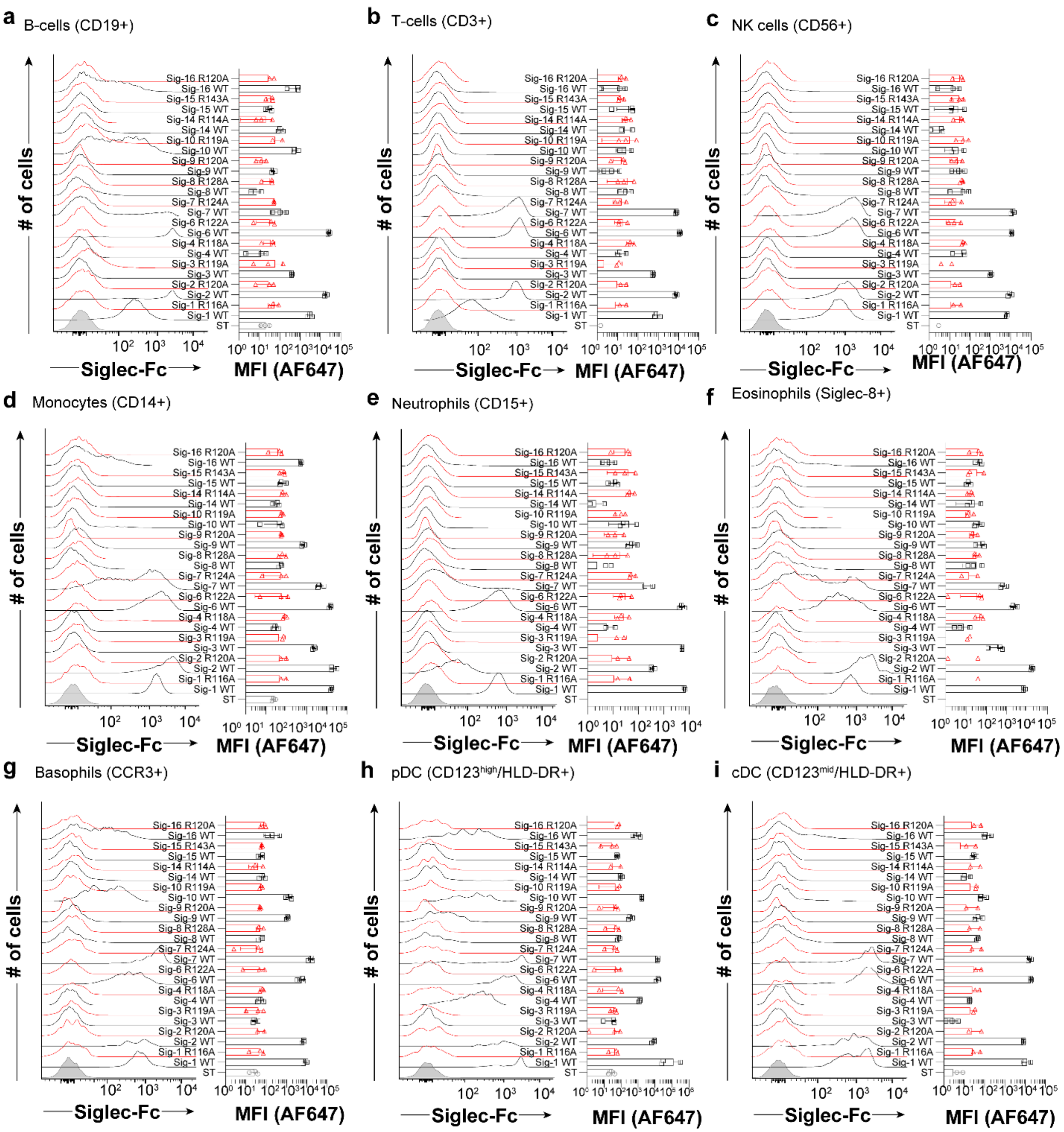


**Supplementary Figure 10: Siglec ligands on human peripheral blood cells.** Flow cytometry data of Siglec-Fc constructs that bound after pre-complexing with Strep-Tactin-AF647 to: **a**, B cells ($CD19^+$); **b**, T cells ($CD3^+$); **c**, NK cells ($CD56^+$); **d**, monocytes ($CD14^+$); **e** neutrophils ($CD15^+$); **f**, eosinophils (Siglec-$8^+$); and **g**, basophils (CCR3+). Each point represents one healthy control subject. Error bars represent +/- standard deviation of three replicates.

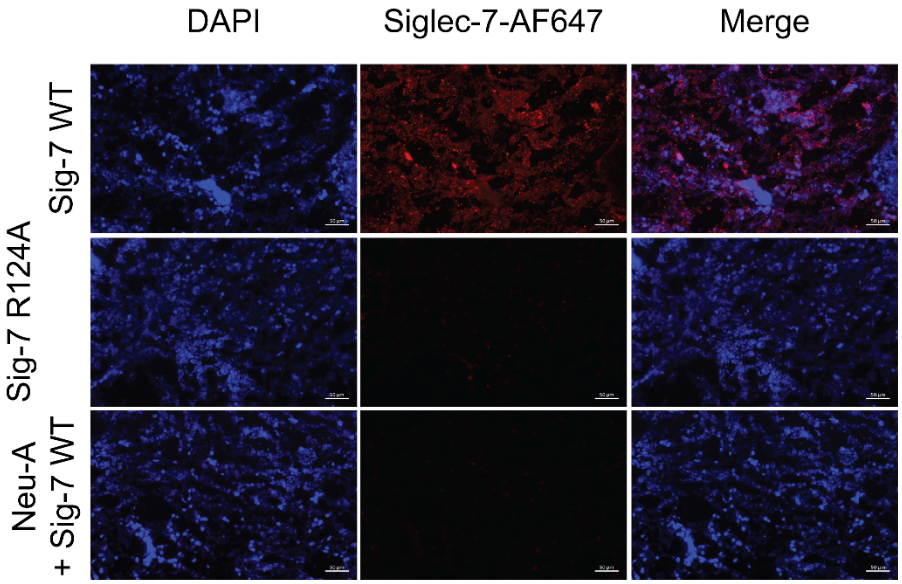


**Supplementary Figure 11: Immunofluorescence staining of human spleen.** Spleen was stained with either Siglec-7-Fc, Siglec-7-Fc R124A, or pre-treated with NeuA and then stained with Siglec-7-Fc WT.

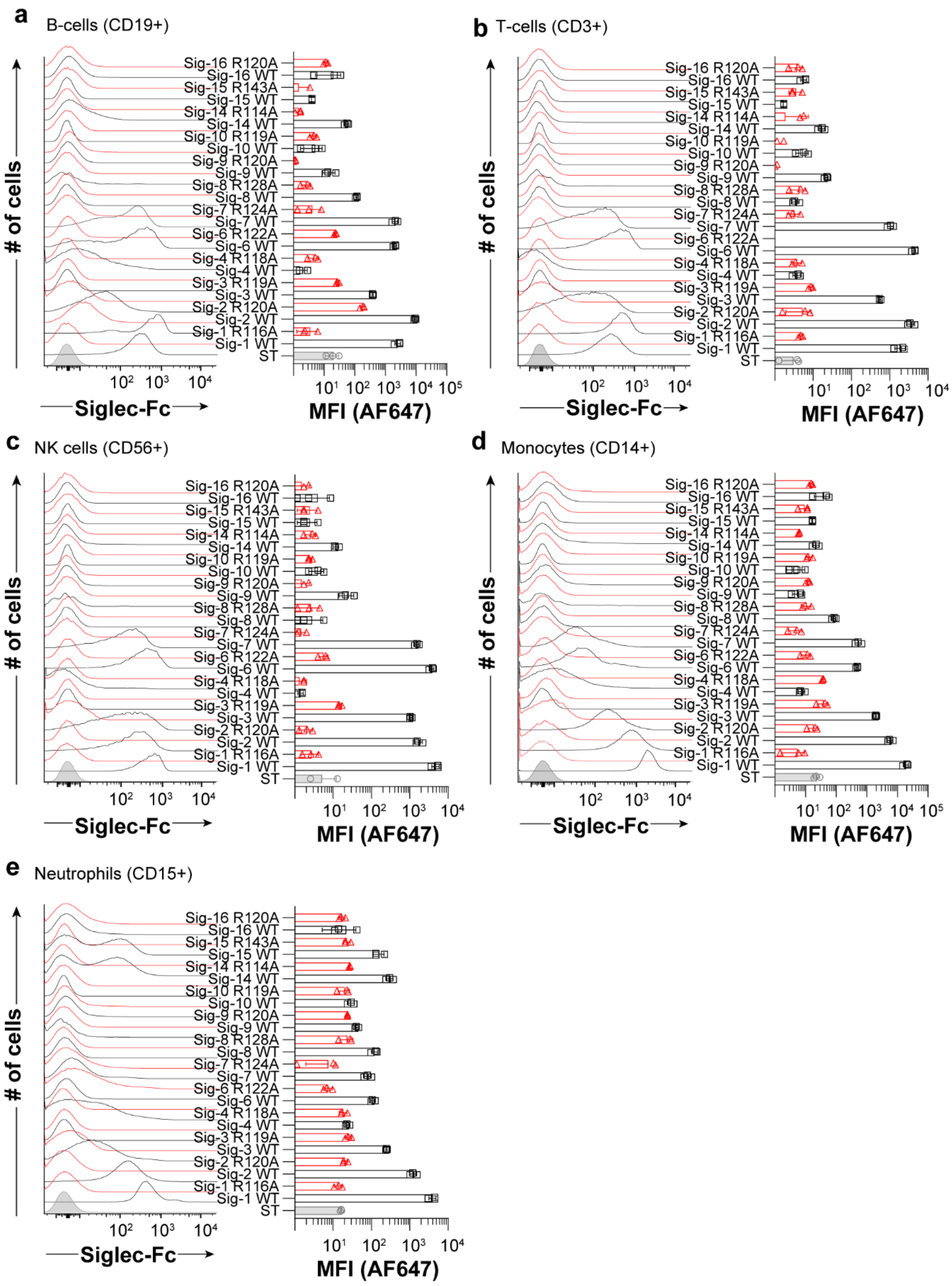


**Supplementary Figure 12: Siglec ligands on human spleen cells.** Flow cytometry data of Siglec-Fc constructs that bound after pre-complexing with Strep-Tactin-AF647 to: **a**, B-cells ($CD19^+$); **b**, T-cells ($CD3^+$); **c**, NK cells ($CD56^+$); **d**, monocytes ($CD14^+$); **e** neutrophils ($CD15^+$). Error bars represent +/- standard deviation of three replicates.

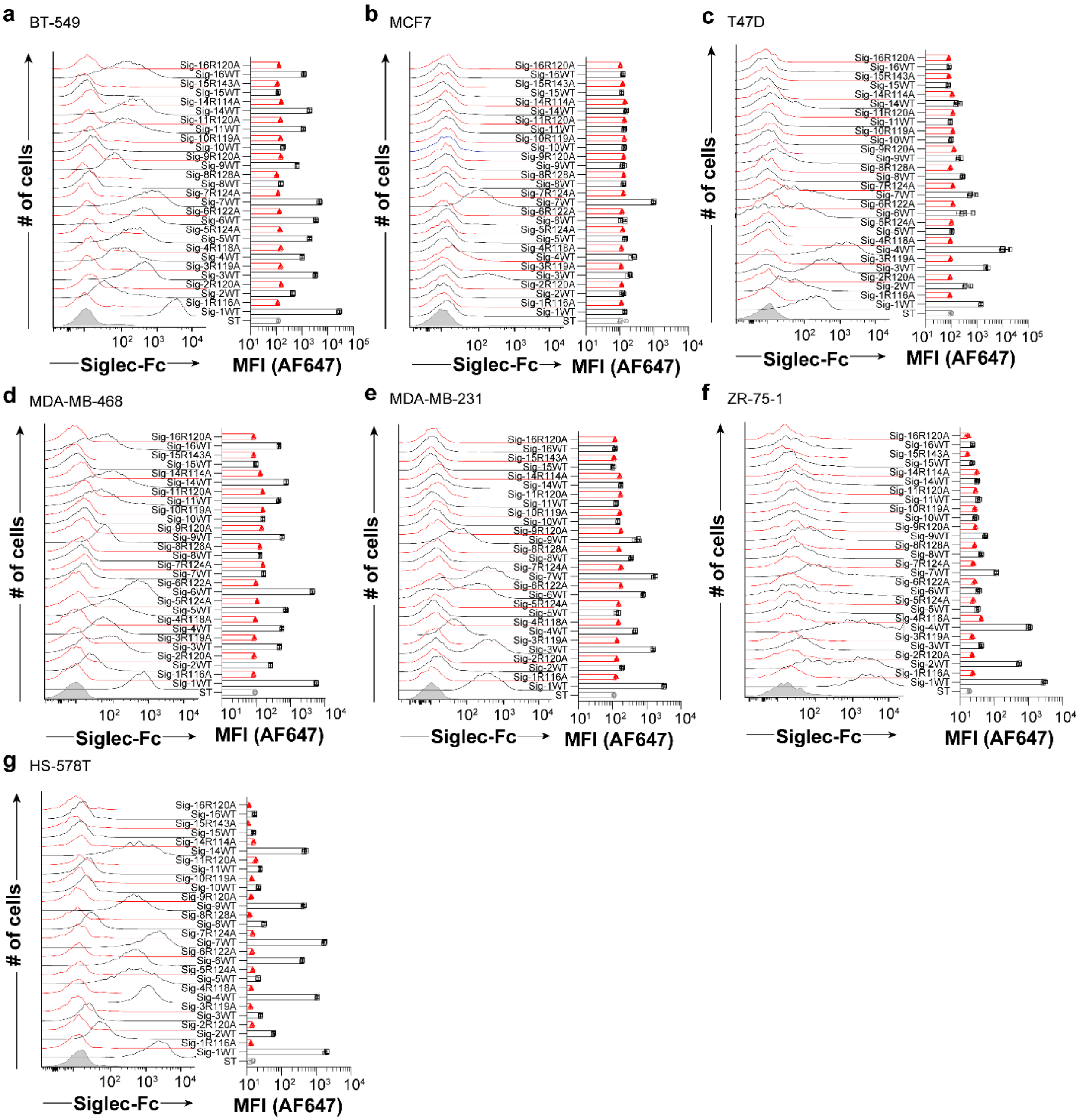

**Supplementary Figure 13: Probing Siglec ligands on breast cancer cell lines.** Flow cytometry data of Siglec-Fc constructs that bound after pre-complexing with Strep-Tactin-AF647 to: **a**, BT549; **b**, MCF7; **c** T47D; **d**, MDA-MB-468; **e**, MDA-MB-231; **f**, ZR-75-1; and **g**, HS-578T cells. Error bars represent +/- standard deviation of three replicates.

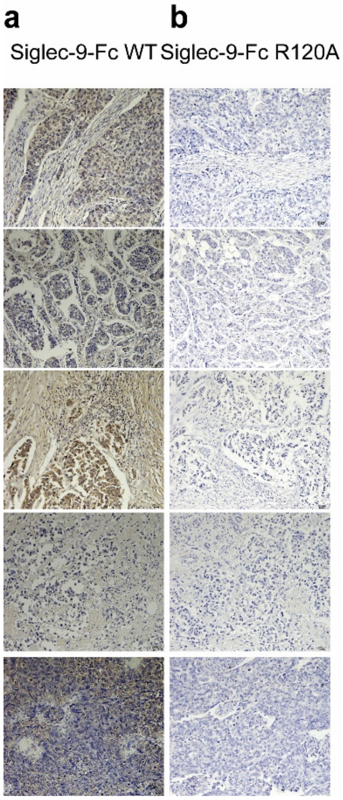


**Supplementary Figure 14:** Immunohistochemistry staining of breast cancer tissues from five different patients. Tissues were stained with Siglec-9-Fc WT (**a**) or Siglec-9-Fc R120A (**b**).

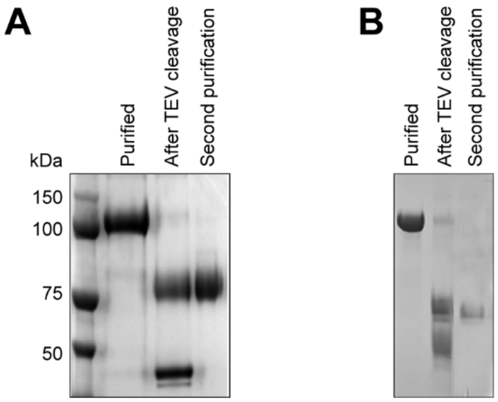


**Supplementary Figure 15: SDS-PAGE gels of the Siglec-Fc construct before and after TEV protease digestion and cleanup by a Nickel column. a**, CD22 and **b**, CD33

**a**

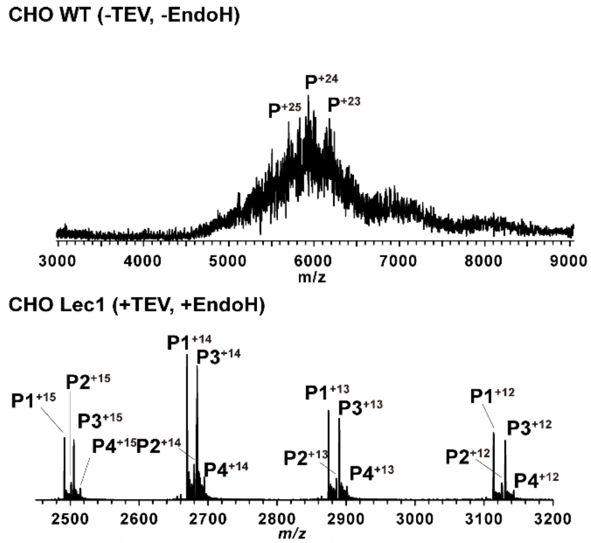


| Protein | Calculated MW (Da) | Experimental MW (Da) |
|---|---|---|
| P1 | 37356.8 | 37356.6 +/- 0.2 |
| P2 | 37502.2 | 37502.2 +/- 0.2 |
| P3 | 37560.0 | 37559.8 +/- 0.2 |
| P4 | 37706.0 | 37705.5 +/- 0.2 |

**b**

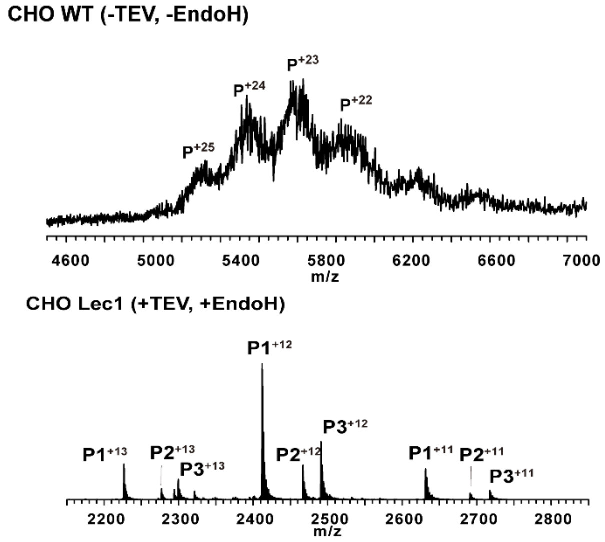


| Protein | Calculated MW (Da) | Experimental MW (Da) |
|---|---|---|
| P1 | 28936.4 | 28936.1 +/- 0.5 |
| P2 | 29593.4 | 29592.6 +/- 0.6 |
| P3 | 29883.4 | 29883.4 +/- 0.6 |

**Supplementary Figure 16: ESI mass spectra acquired** in positive mode for aqueous ammonium acetate (200 mM, pH 7.0) solutions of WT CHO full Fc-chimera and Lec-1 CHO cells following digestion with TEV, and after further treatment with Endo-H and their associated calculated and experimental molecular weight for: **a**, CD22 and **b**, CD33

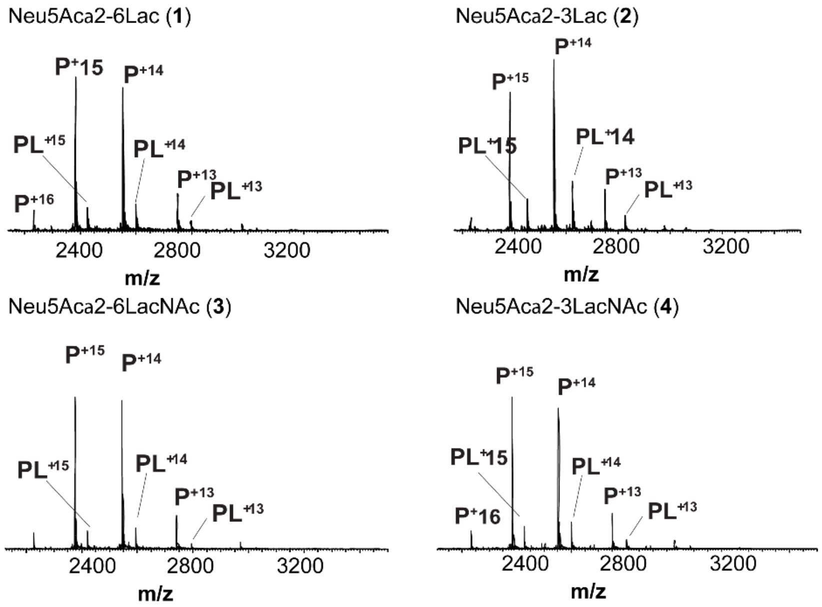


**Supplementary Figure 17: ESI mass spectra data for Siglec-1 binding to four different trisaccharadies.** ESI mass spectra were acquired in positive mode for aqueous ammonium acetate (200 mM, pH 7.0 and 25 °C) solutions of 3.6 μM Siglec-1 (P = Siglec1 fragment) and 80 μM of Neu5Acα2-6Lac (**1**), Neu5Acα2-3Lac (**2**), Neu5Acα2-6LacNAc (**3**), and Neu5Acα2-3LacNAc (**4**). Cytochrome C (1 μM), which served as $P_{ref}$, was added to each solution to correct for nonspecific binding.

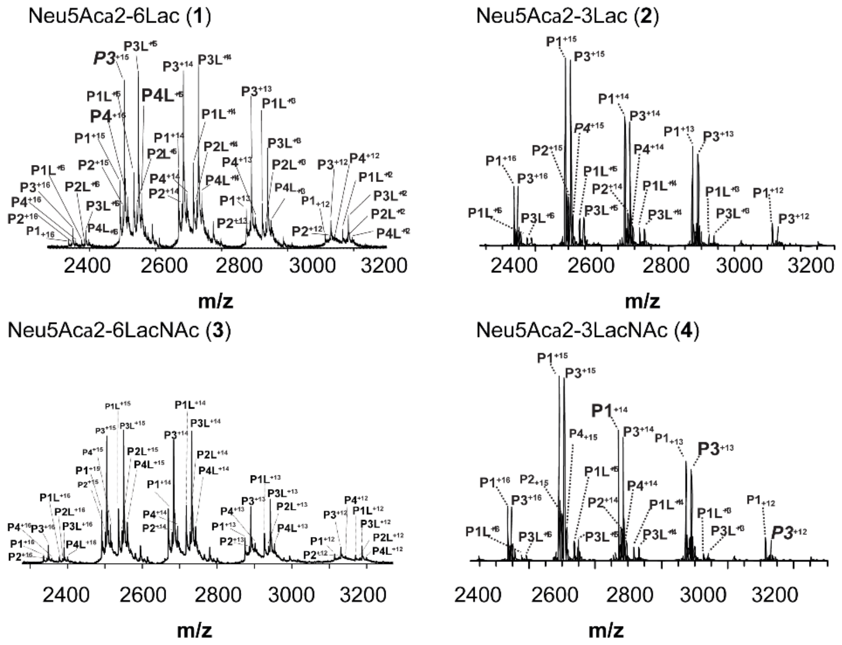


**Supplementary Figure 18: ESI mass spectra data for CD22 binding to four different trisaccharadies.** ESI mass spectra acquired in positive mode for aqueous ammonium acetate (200 mM, pH 7.0 and 25 °C) solutions of 20 μM CD22 (P = CD22 fragment), and 80 μM of Neu5Acα2-6Lac (**1**), Neu5Acα2-3Lac (**2**), Neu5Acα2-6LacNAc (**3**), and Neu5Acα2-3LacNAc (**4**). Cytochrome C (1 μM), which served as $P_{ref}$, was added to each solution to correct for nonspecific binding.

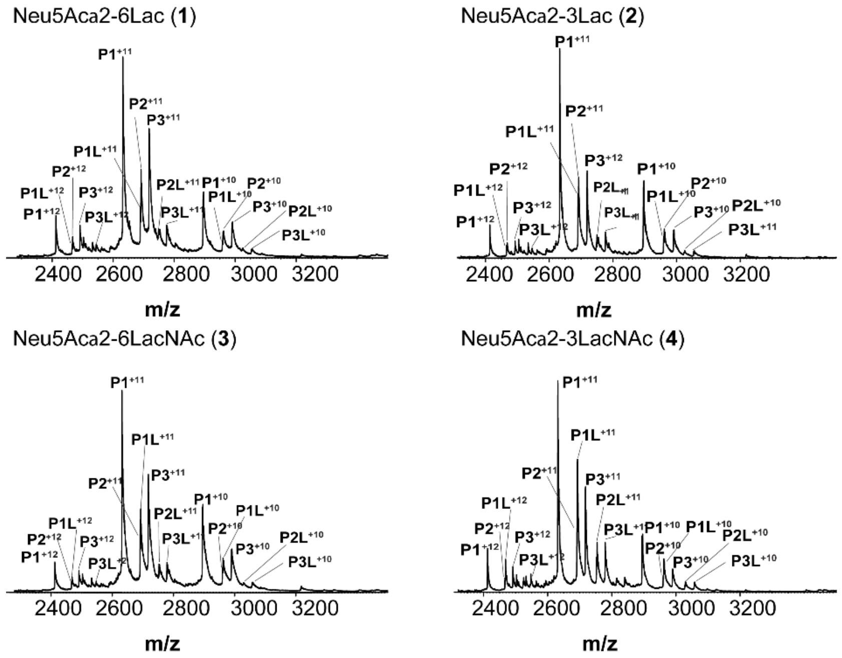


**Supplementary Figure 19: ESI mass spectra data for CD33 binding to four different trisaccharadies.** ESI mass spectra acquired in positive mode for aqueous ammonium acetate (200 mM, pH 7.0 and 25 °C) solutions of 5.3 μM CD33 (P = CD33 fragment), and 320 μM of Neu5Acα2-6Lac (**1**), Neu5Acα2-3Lac (**2**), Neu5Acα2-6LacNAc (**3**), and Neu5Acα2-3LacNAc (**4**). Cytochrome C (1 μM), which served as $P_{ref}$, was added to each solution to correct for nonspecific binding.

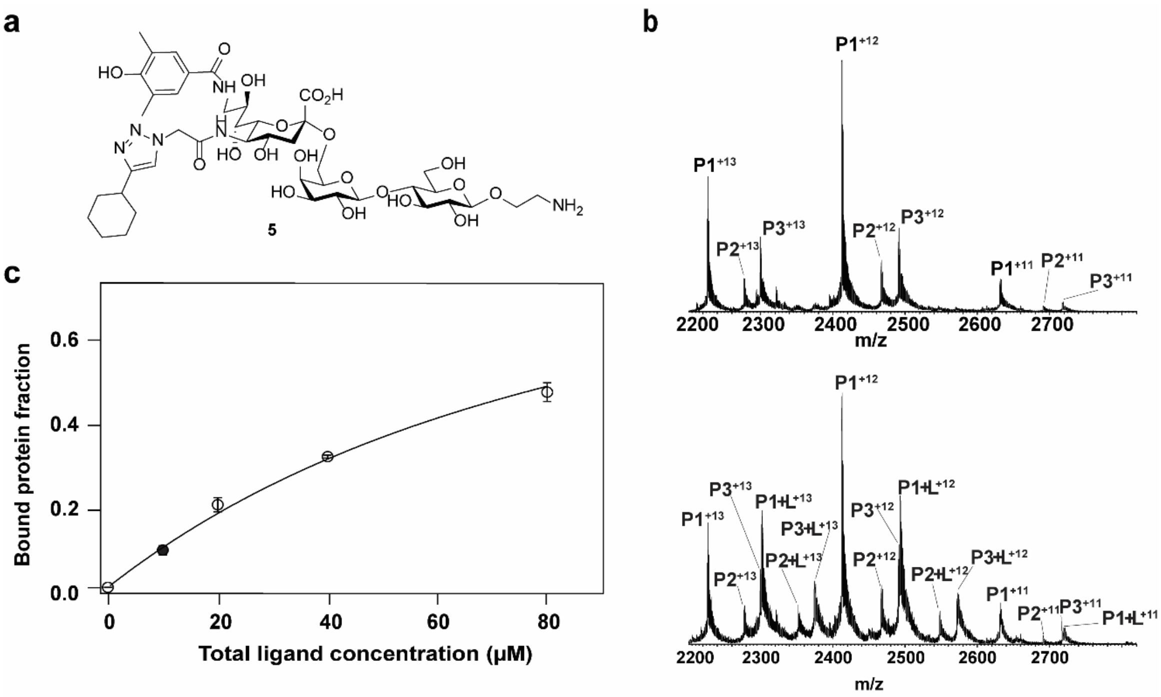


**Supplementary Figure 20: ESI mass spectra data for CD33 binding to high-affinity CD33L.** **a**, Structure of 5, 9-bifunctional-6-sialyllactose, the high-affinity CD33 ligand (CD33L, compound **5**). **b**, Mass spectrum of 4.2 μM of CD33 (+TEV, +Endo-H) before incubation (top), and after incubation with 50 μM of CD33L (bottom). **c**, Fraction of the ligand-bound CD33 plotted versus initial concentration of CD33 measured by ESI-MS.

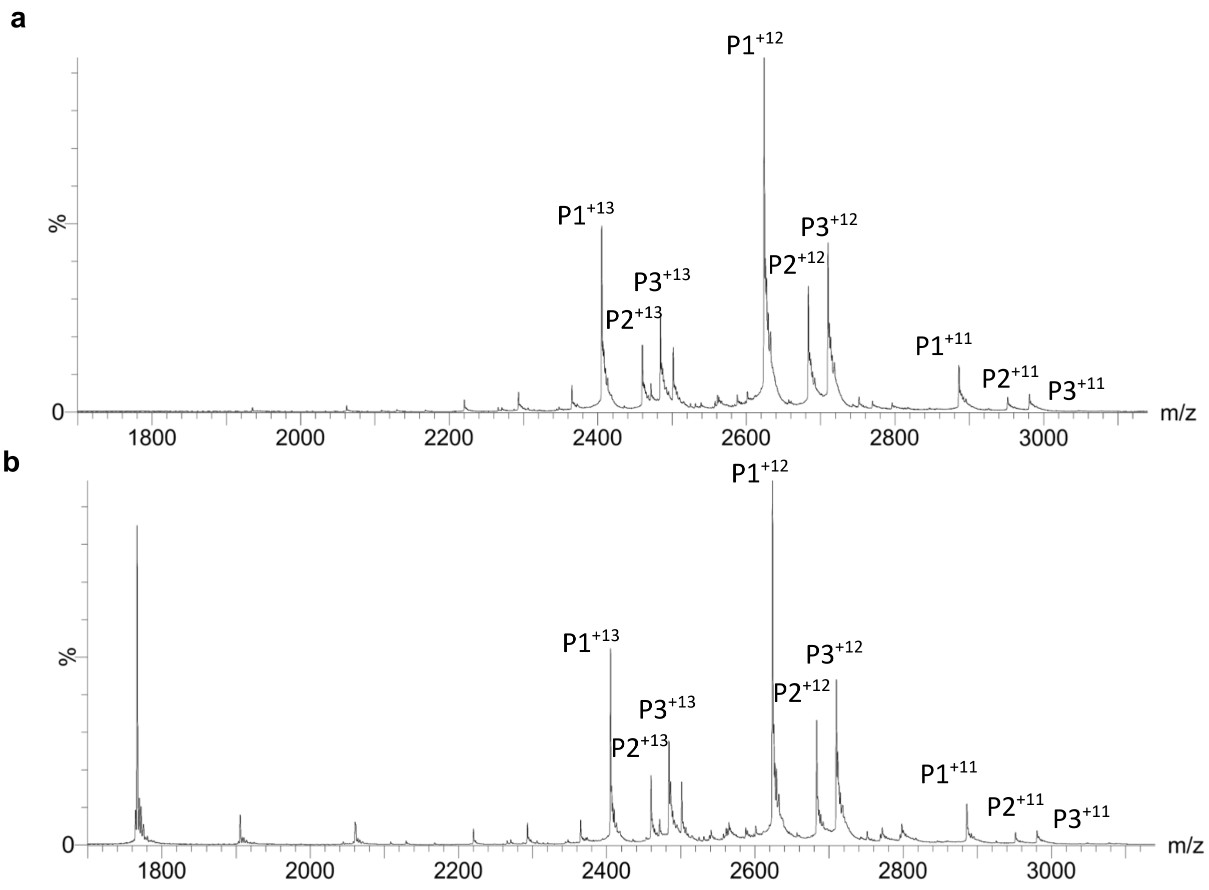


**Supplementary Figure 21: The R119A mutant of CD33 does not bind the high-affinity CD33L.** Mass spectrum of 4.2 µM of CD33 R119A (+TEV, +Endo-H) without (**a**), and with 50 µM of CD33L (compound **5**) (**b**).

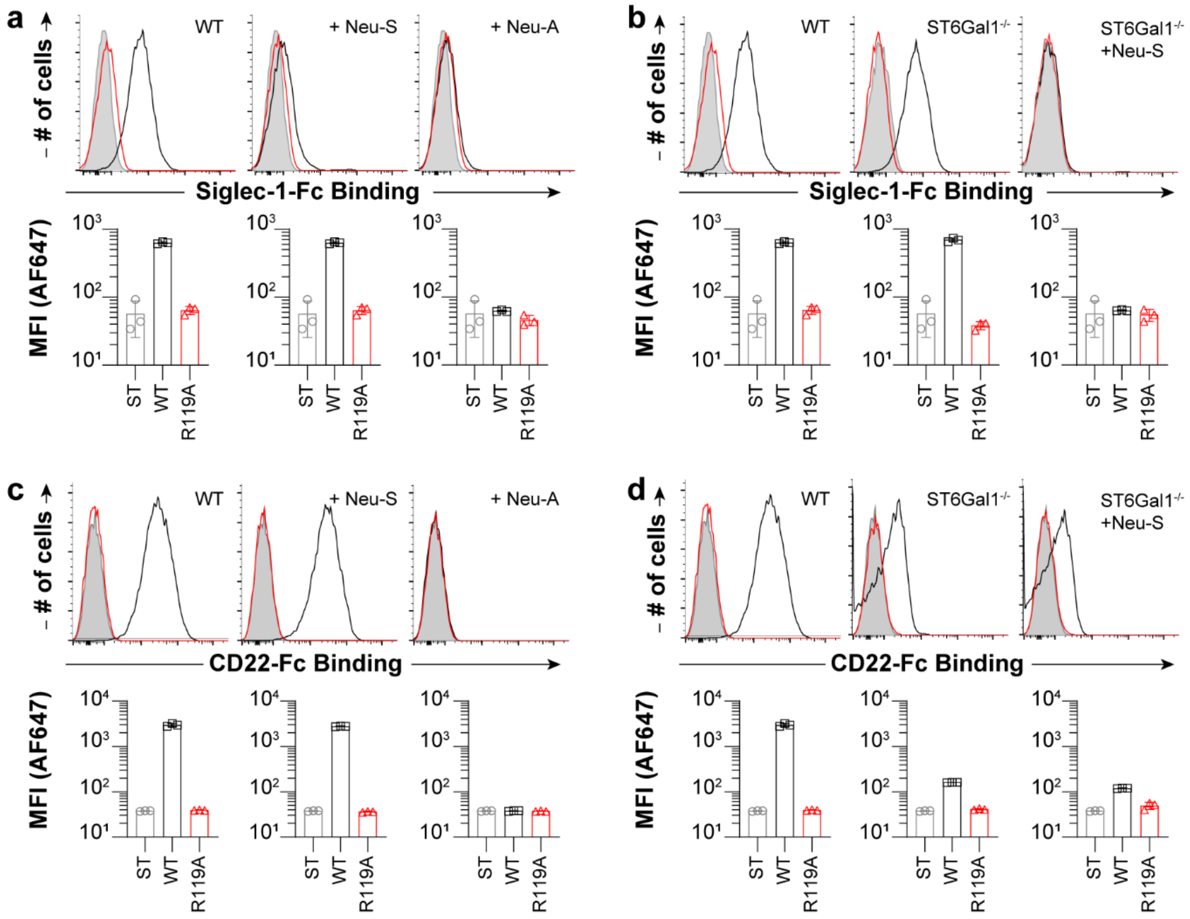


**Supplementary Figure 22: Cellular glycan ligands of Siglec-1 and CD22. a**, Staining of U937 cells treated with an α2-3-specific neuraminidase (Neu-S) or broadly acting neuraminidase (Neu-A) with Siglec-1-Fc pre-complexed with Strep-Tactin-AF647. **b**, Staining of WT, ST6Gal1$^{-/-}$ U937 cells, and ST6Gal1$^{-/-}$U937 cells treated with Neu-S with Siglec-1-Fc pre-complexed with Strep-Tactin-AF647. **c**, Staining of U937 cells treated with Neu-S or Neu-A with CD22-Fc pre-complexed with Strep-Tactin-AF647. **d**, Staining of WT, ST6Gal1$^{-/-}$ U937 cells, and ST6Gal1$^{-/-}$U937 cells treated with Neu-S with CD22-Fc pre-complexed with Strep-Tactin-AF647.

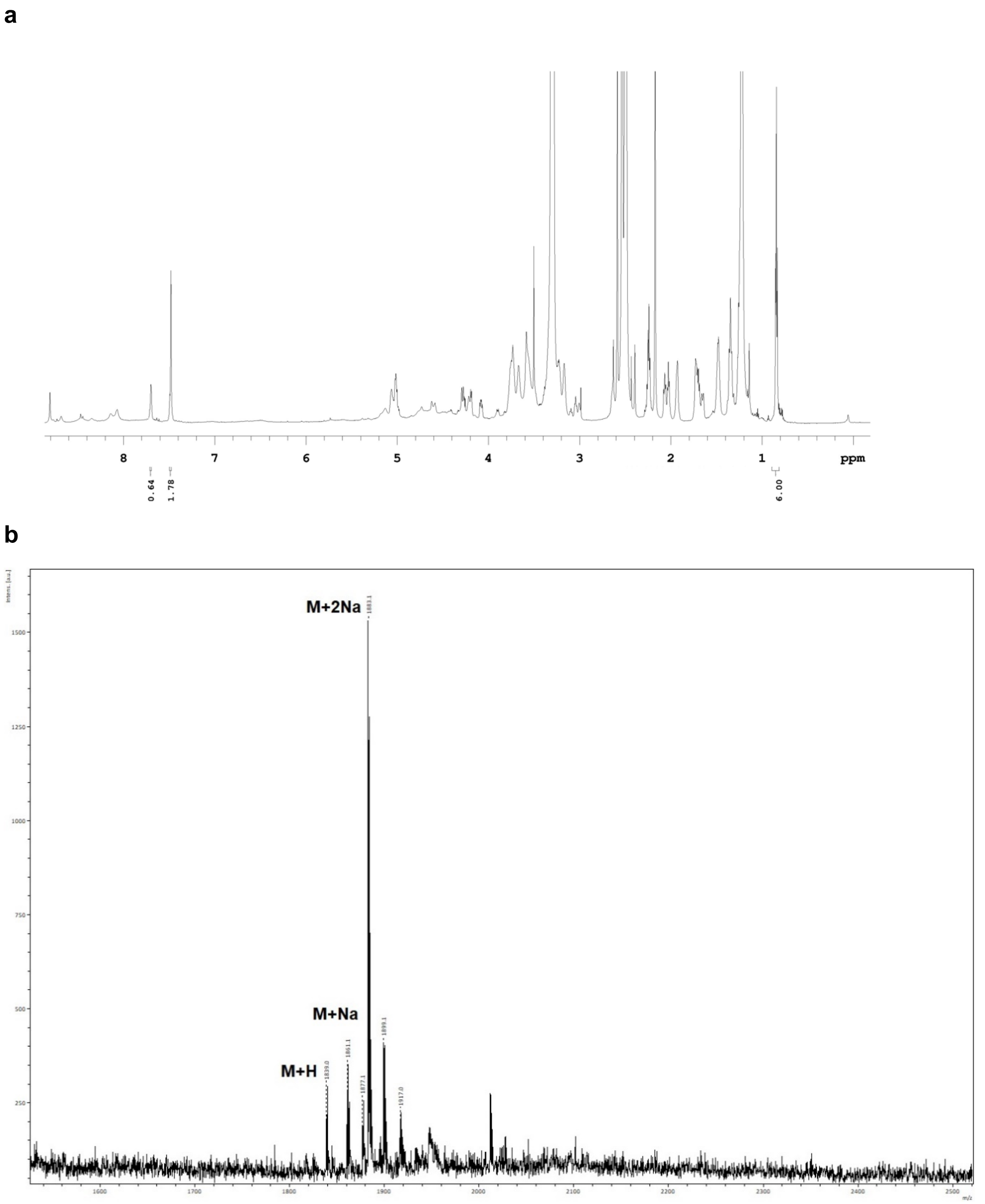


**Supplementary Figure 23: Analytical analysis of CD33L-DSPE. a**, $^1$H-NMR in DMSO-$D_6$, 700 MHz. **b**, MALDI-TOF mass spectrometry of CD33L-DSPE lipid conjugate.

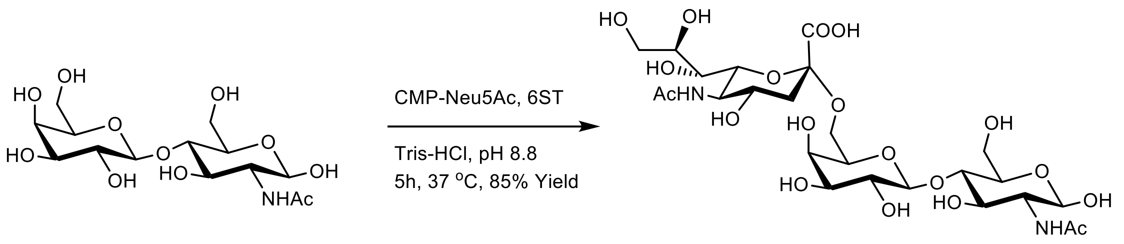


**1**

**Supplementary Scheme 1: Synthesis of *N*-acetylneuraminosylα2-6-*O*-D-galactopyranosylβ1-4-*O*-D-glucopyranose (Neu5Acα2-6Lac,** compound **1).**

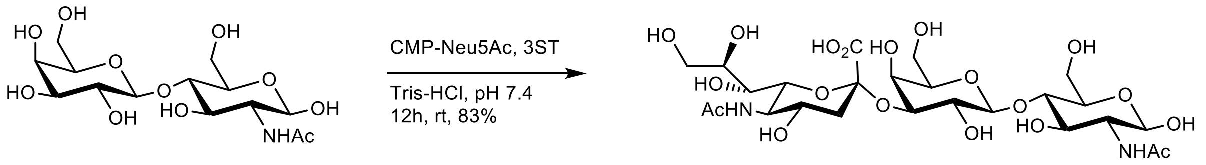


**2**

**Supplementary Scheme 2: Synthesis of *N*-acetylneuraminosylα2-3-*O*-D-galactopyranosylβ1-4-*O*-D-glucopyranose (Neu5Acα2-3Lac,** compound 2**).**

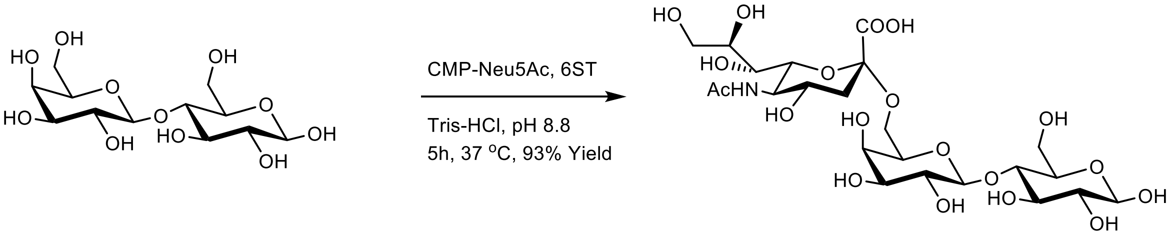


**3**

**Supplementary Scheme 3: Synthesis of *N*-acetylneuraminosylα2-6-*O*-D-galactopyranosylβ(*1-4*)-2-(acetylamino)-2-deoxy-D-glucopyranose (Neu5Acα2-6LacNAc,** compound **3).**

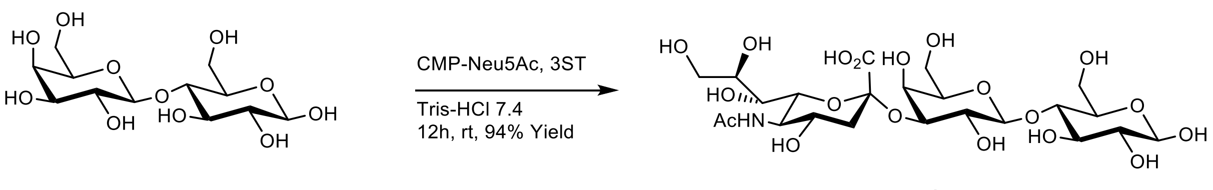


**4**

**Supplementary Scheme 4: Synthesis of *N*-acetylneuraminosylα2-3-*O*-D-galactopyranosylβ(*1-4*)-2-(acetylamino)-2-deoxy-D-glucopyranose (Neu5Acα2-3LacNAc,** compound **4).**

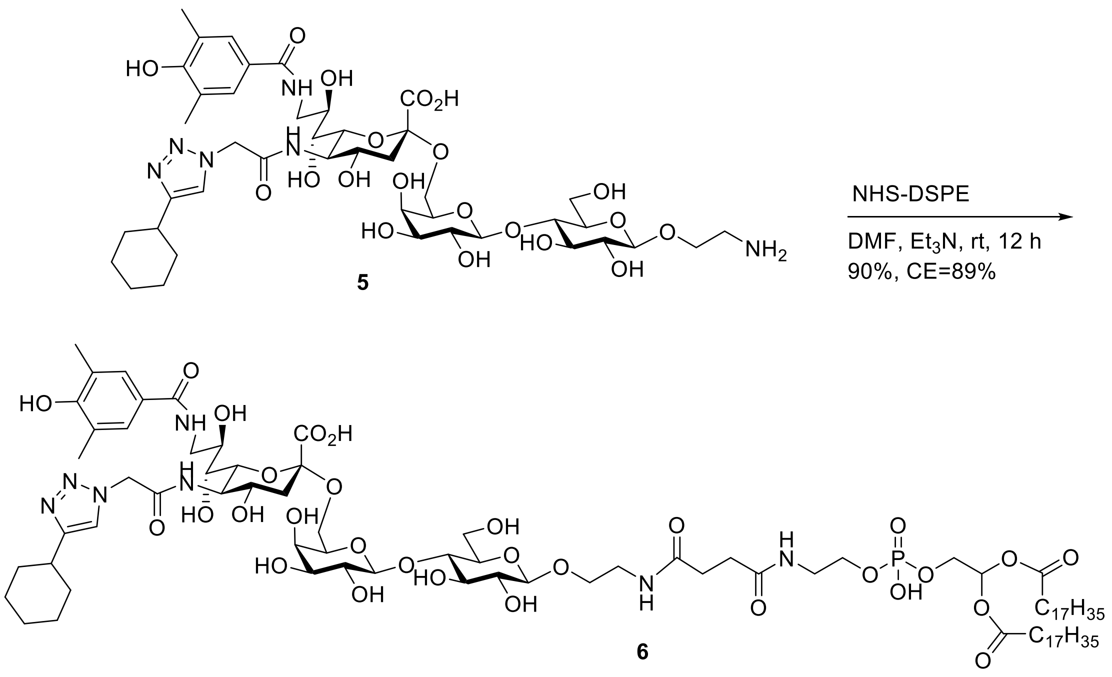


**Supplementary Scheme 5: Synthesis of CD33L-DSPE lipid conjugate (**compound **6).**
